# Rac rather than Rho drives the early transcriptional response to extracellular matrix stiffness and mediates repression of ATF3

**DOI:** 10.1101/2022.04.27.489717

**Authors:** Irène Dang, Yongho Bae, Joseph A. Brazzo, Richard K Assoian

## Abstract

The Rho family GTPases, Rac and Rho, play critical roles in transmitting mechanical information contained within the extracellular matrix (ECM) to the cell. Rac and Rho have well described roles in regulating stiffness-dependent actin remodeling, proliferation and motility. However, much less is known about the relative roles of these GTPases in stiffness-dependent transcription, particularly at the genome-wide level. Here, we selectively inhibited Rac and Rho in mouse embryonic fibroblasts cultured on deformable substrata and used RNA sequencing to elucidate and compare the contribution of these GTPases to the early transcriptional response to ECM stiffness. Surprisingly, we found that Rac activation is almost exclusively responsible for the initial transcriptional response to ECM stiffness. We also identified Activating Transcription Factor 3 (ATF3) as a major target of stiffness/Rac signaling and show that ATF3 repression by ECM stiffness connects the stiffness-dependent activation of Rac to the induction of cyclin D1.

## INTRODUCTION

Cell adhesion to its surrounding extracellular matrix (ECM) is essential for viability, motility and proliferation of many cell types. The ECM is complex and composed of both protein and non-proteinaceous components. Adhesion to the ECM provides chemical information to the cell because many distinct ECM components bind to specific receptors on the cell surface to regulate intracellular signaling, often in concert with growth factor/cytokine-mediated activation of receptor tyrosine kinases (Boudreau and Jones, 1999; Assoian and Schwartz, 2001; Larsen *et al*., 2006). Additionally, more recent work has shown that the ECM also provides mechanical information to the cell, and the rigidity or stiffness of the ECM affects many of the signaling events and cellular fates originally attributed to cell adhesion (Discher *et al*., 2005; Suresh, 2007; Janmey *et al*., 2015). Cells can sense the stiffness of their microenvironment when ECM proteins such as the fibrillar collagens bind to their specific cell surface receptors in the integrin family (Lee and Juliano, 2004; Bourgot *et al*., 2020; Tang, 2020; Sun, 2021). Other ECM proteins such as fibronectin (FN) and vitronectin also bind to specific integrins and contribute to the information content of the cell’s microenvironment.

The cytoplasmic domain of integrins lacks intrinsic kinase activity, but they associate directly and indirectly with a number of signaling molecules in dynamic macromolecular structures called focal complexes (FCs) and focal adhesions (FAs). FCs are near the cell periphery and unstable but mature into FAs under force (Geiger and Bershadsky, 2001; Geiger and Yamada, 2011; Burridge and Wittchen, 2013; Burridge and Guilluy, 2016; Wolfenson *et al*., 2018). FAs contain many signaling molecules (Sastry and Burridge, 2000; Zaidel-Bar *et al*., 2007; Horton *et al*., 2016) and are also non-covalently linked to actin stress fibers, which mediate actomyosin contraction, generate intracellular tension, and reinforce FA maturation and stabilization (Geiger and Bershadsky, 2001; Geiger and Yamada, 2011; Lessey *et al*., 2012; Burridge and Wittchen, 2013; Burridge and Guilluy, 2016; Wolfenson *et al*., 2018). Thus, increased ECM stiffness promotes FA formation and regulates cytoplasmic signaling pathways that control mechanotransduction to the nucleus and a number of stiffness-sensitive cell fates including motility, differentiation, proliferation and transformation (Paszek *et al*., 2005; Yeung *et al*., 2005; Engler *et al*., 2006; Assoian and Klein, 2008; Zajac and Discher, 2008; Janmey *et al*., 2015; Wei *et al*., 2015; Talwar *et al*., 2021).

The Rac and Rho GTPases have pleiotropic effects on cells and tissues and are often involved in mechanosensitive signaling pathways and influence the cell fates described above (Zohn *et al*., 1998; Sahai and Marshall, 2003; Burridge and Wennerberg, 2004; Hall, 2005; Provenzano and Keely, 2011; Lawson and Burridge, 2014; Pasapera *et al*., 2014; Ridley, 2015). Rac activation is critical for FC formation and cell spreading while Rho activity has an established role in stress fiber and FA formation and enforcement (Zohn *et al*., 1998; Sahai and Marshall, 2003; Burridge and Wennerberg, 2004; Hall, 2005; Provenzano and Keely, 2011; Lawson and Burridge, 2014; Pasapera *et al*., 2014; Ridley, 2015). Both Rac and Rho play important roles in cell motility (Sahai and Marshall, 2003; Raftopoulou and Hall, 2004; Hall, 2005; Lawson and Burridge, 2014; Ridley, 2015). In contrast, we have previously reported that the activation of Rac, but not Rho, is required for stimulatory effects of ECM stiffness in G1 phase cell cycle progression (Klein *et al*., 2009; Bae *et al*., 2014). In this signaling pathway, the stiffness-dependent activation of FAK within FAs leads to the activation of p130Cas and eventually DOCK180, a Rac guanine nucleotide exchange factor. The consequent stiffness-dependent activation of Rac and induction of lamellipodin is required for the mid-G1 phase expression of cyclin D1 (Klein *et al*., 2009; Bae *et al*., 2014; Brazzo *et al*., 2021). Among its effects, cyclin D1 is perhaps best understood as an activator of cyclin-dependent kinase 4/6, which in turn, contributes to the release of E2Fs from the retinoblastoma protein family and promotes S phase entry (Sherr, 1993, 1995; Resnitzky and Reed, 1995; Weinberg, 1995). Thus, the stiffness-dependent activation of Rac plays an important role in the stimulatory effect of ECM stiffness on cell cycling. The third Rho family GTPase, Cdc42, is responsible for filopodia formation and directional motility (Cerione, 2004; Raftopoulou and Hall, 2004; Hall, 2005; Ridley, 2006), but its role in mechano-sensitive cell fates is not as well understood.

In addition to their effects on cytoplasmic signaling, both Rac and Rho have been reported to regulate transcription. The transcriptional effect is best understood for Rho, as stimulatory effects of Rho on actin polymerization lead to nuclear translocation of the MAL (also referred to as myocardin-related transcription factors and megakaryoblastic leukemia-1 and −2) co-activator of SRF genes (Sotiropoulos *et al*., 1999; Miralles *et al*., 2003; Pipes *et al*., 2006; Parmacek, 2007; Zhao *et al*., 2007). Additionally, both Rho and Rac have been implicated in the nuclear translocation of the YAP/Taz transcriptional coactivators (Dupont *et al*., 2011; Aragona *et al*.,2013; Jang *et al*., 2017; Talwar *et al*., 2021). Nevertheless, much remains unknown about the relative contributions of Rac and Rho to the transcriptional response to ECM stiffness, particularly at the genome-wide scale. To address this gap in understanding, we used next generation sequencing to determine and compare the effects of Rac versus Rho on the early transcriptional response to ECM stiffness. Despite the fact that Rac and Rho have similar effects on intracellular stiffness, we find that Rac is almost completely responsible for the initial transcriptional response to ECM stiffness. Additionally, we show that the levels of Activating Transcription Factor 3 (ATF3) are repressed by both high ECM stiffness and increased Rac activity. Stiffness-mediated ATF3 repression links mechanosensitive activation of Rac to cyclin D1 gene expression and S phase entry.

## RESULTS

### Rac and Rho similarly affect intracellular stiffness during early stages of cell adhesion and spreading

To compare the effects of Rac and Rho on the early transcriptional response to ECM stiffness, we treated serum-starved mouse embryo fibroblasts (MEFs) with drugs that acutely and specifically inhibit either Rac (EHT1864, a potent Rac family GTPase inhibitor) or Rho (CT04, cell permeable Rho inhibitor). We plated these drug-treated cells on fibronectin (FN)-coated polyacrylamide hydrogels in the presence of fetal bovine serum (FBS), using a hydrogel stiffness that mimics the stiffness found at sites of proliferation *in vivo* (20-25 kPa; hereafter called “stiff”) (Klein *et al*., 2009). This stiffness range also promotes motility and S-phase entry in cultured cells (Klein *et al*., 2009; Bae *et al*., 2014; Razinia *et al*., 2017; Shutova *et al*., 2017). We also used a lower stiffness (4-6 kPa; hereafter called “soft”) that failed to stimulate proliferation and motility (Klein *et al*., 2009; Bae *et al*., 2014; Razinia *et al*., 2017; Shutova *et al*., 2017). MEFs were cultured for 1 h before collection and analysis. Using this approach, we tested EHT1864 and CT04 for their ability to inhibit the stiffness-dependent GTP-loading of Rac and Rho, respectively. We identified concentrations of these drugs that resulted in strong inhibition of the respective GTPase target without significant effect on the other Rho family GTPases (Fig. 1A). F-actin abundance was reduced in response to both EHT1864 and CT04 (Fig. 1B-C). In addition, Rac, but not Rho, inhibition reduced cell area (Fig. 1B and 1D), a consequence of impaired cell spreading (Fig. S1). Importantly, both Rac and Rho inhibition similarly reduced intracellular stiffness as measured by atomic force microscopy (Fig. 1E).

**Figure 1.**
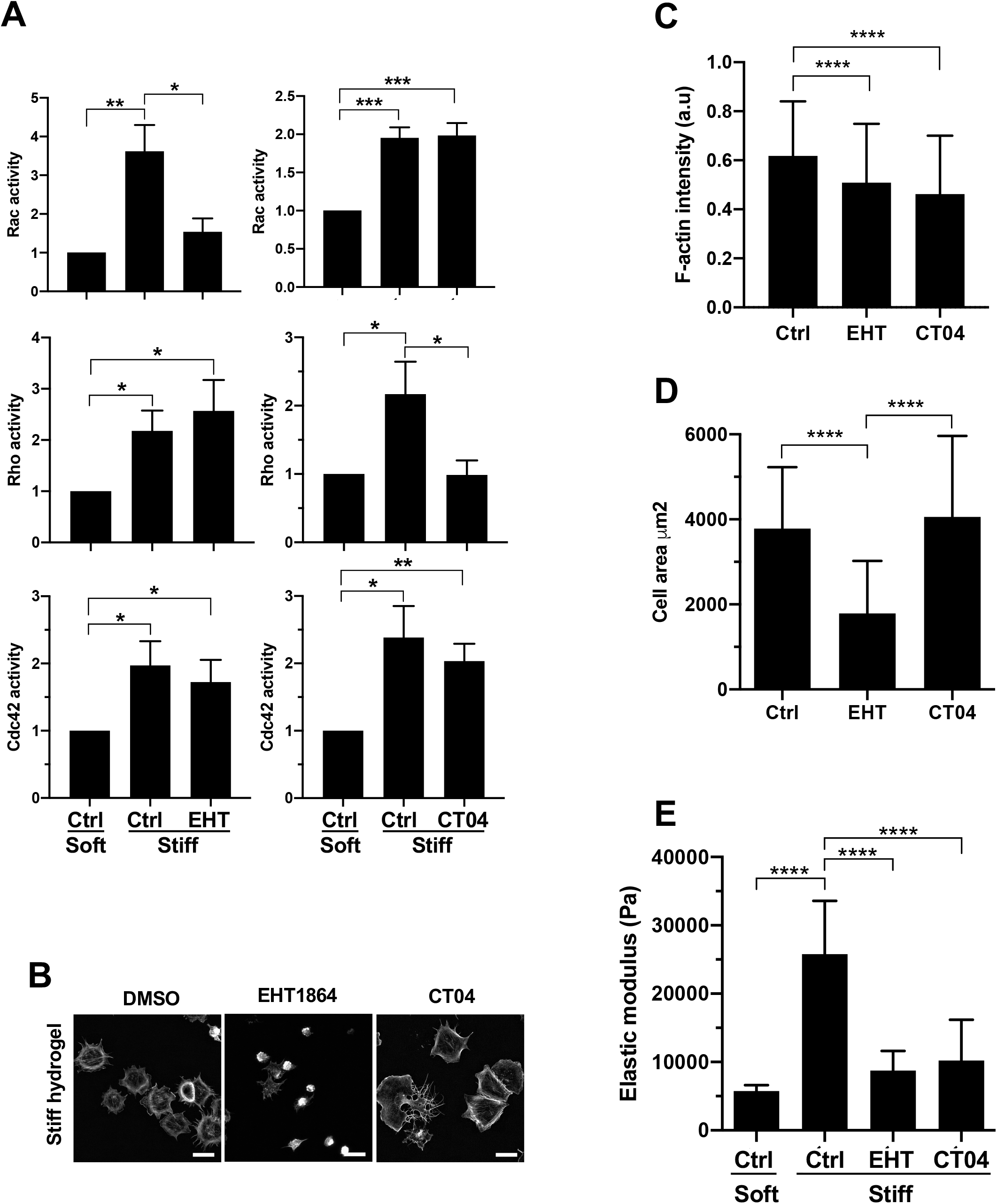
Rac and Rho inhibition similarly reduce f-actin staining cell stiffness. Serum-starved MEFs pre-incubated with CT04 (Rho inhibitor) or EHT1864 (Rac inhibitor) were plated on FN-coated hydrogels and stimulated with 10% FBS for 1 h. **(A)** Rho-family GTPase activity was measured by G-LISA and plotted relative to the activity on the soft hydrogels (n=4). Graphs show mean + SEM. **(B)** Maximum projection images of phalloidin-stained MEFs taken by confocal microscopy. Scale bar = 50 μm. **(C)** Quantification of phalloidin-stained MEFs was measured and normalized to cell area with Image J. Results show mean and SD; n=137 (DMSO), 165 (EHT1864), and 152 (CT04) cells accrued from 3 independent experiments. **(D)** Cell areas were measured with ImageJ. Results show mean and SD; n=259 (DMSO), 308 (EHT1864), and 222 (CT04) cells accrued from 3 independent experiments. **(E)** Cell stiffness was determined by AFM. The graph shows mean + SEM (n=34 cells per condition accrued over 4 independent experiments).

### Rac dominates the early transcriptional response to ECM stiffness and mediates the stiffness-dependent repression of ATF3

Using the experimental conditions identified in Fig. 1, we performed an RNA sequencing (RNASeq) to interrogate the relative roles of Rac and Rho in transducing the early transcriptional response to ECM stiffness. The results showed that of 483 genes stimulated by ECM stiffness, one third (158 genes) were inhibited by EHT1864, but only 1% (6 genes) were inhibited by CT04 (Fig. 2A and Table S1). Similarly, Rac was dominant over Rho when examining the genes that were inhibited by ECM stiffness (Fig. 2B and Table S2): about 40% of stiffness-inhibited genes showed an increased abundance in response to EHT1864 (123 of 327) but only 6% showed increased abundance in response to CT04 (19 of 327). Very few stiffness-regulated genes were affected by both the Rac and Rho inhibitors (Fig. 2A-B and Tables S1 and S2). Moreover, Fisher exact tests showed that the overlap between the stiffness-regulated and EHT1864-regulated genes was significant (p<0.0001 and 0.019 for the stiffness-stimulated and -inhibited, genes respectively) whereas the overlap between stiffness-regulated genes and the few CT04-regulated genes was not significant (p=0.2 and 0.31 for stiffness-stimulated and -inhibited, genes respectively). Thus, Rac rather than Rho drives the early transcriptional program in response to ECM stiffness.

**Figure 2.**
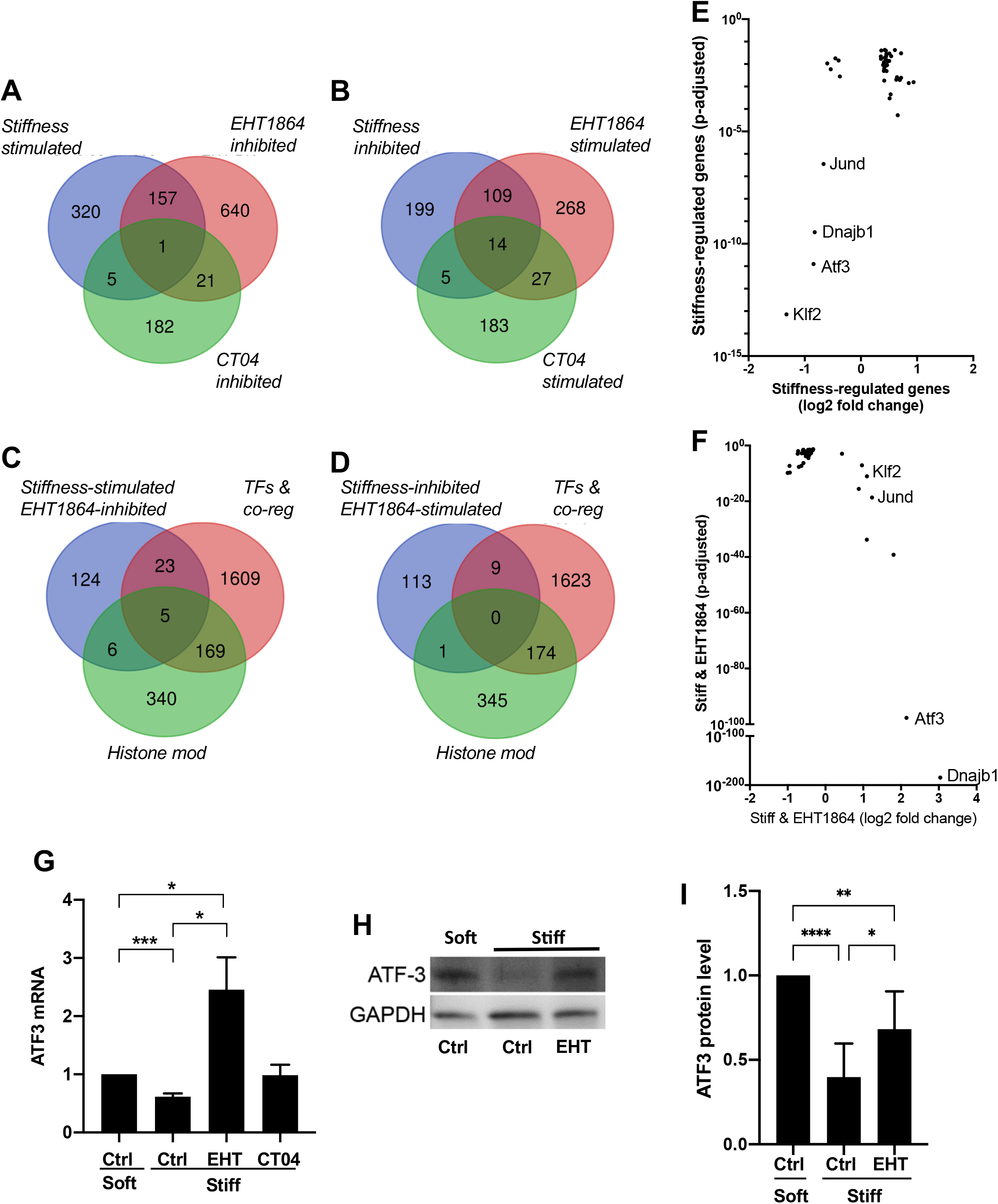
Rac drives the initial genome-wide response to ECM stiffness and preferentially represses ATF3. **(A-B)** Venn diagrams of differentially expressed genes in MEFs cultured with 10% FBS for 1 hour on stiff versus soft hydrogels, stiff hydrogels ± EHT1864, or stiff hydrogels ± CT04. **(C-D)** The genes regulated by ECM and Rac (163 stiffness-stimulated/EHT-inhibited and 123 stiffness-inhibited/EHT-stimulated genes; see Figs. 2A and B, respectively) were compared to GO gene lists for transcription factors, transcription co-regulators and histone modifiers. **(E-F)** Log2 fold-changes and adjusted p-values of the genes regulated by ECM stiffness/Rac and contained within the GO terms described above. **(G)** Serum-starved MEFs were plated on soft or stiff FN-coated hydrogels with 10% FBS for 1 h in the presence of DMSO (Ctrl), EHT1864, or CT04. Total RNA was extracted, and ATF3 mRNA levels were quantified by RT-qPCR. The graph show mean + SEM (n=4). **(H-I)** Serum-starved MEFs were cultured as in panel C but with DMSO or EHT1864. Cell lysates were analyzed by western blot; results from 6 independent experiments were quantified using Image J. The graph shows mean + SD.

Several studies have demonstrated a role for Rho signaling in mechanosensitive transcription, most commonly through MAL or YAP/Taz (see Introduction). The striking absence of Rho signals in our RNASeq data set does not question these findings but rather emphasizes that the *initial* transcriptional response to ECM stiffness is mediated by Rac, not Rho.

In an effort to better understand this genome-wide response, we searched for transcriptional regulators within the set of genes that were inversely regulated by ECM stiffness and Rac. The 158 genes that were stimulated by ECM stiffness and inhibited by EHT1864 (refer to Fig. 2A) and the 123 genes that were inhibited by ECM stiffness and stimulated by EHT1864 (refer to Fig. 2B) were compared to the Gene Ontology (GO) lists of transcription factors, transcriptional coregulators, and histone modifiers (Figs. 2C-D and Table S3). The overlapping genes (34 and 10 in Fig. 2C and 2D, respectively) were then graphed by fold-change and adjusted p-value to visualize the relative robustness of their responses to ECM stiffness and Rac inhibition with EHT1864 (Fig. 2E-F and Table S4). The results showed that, as a group, neither transcription co-regulators nor histone modifiers were strongly regulated during the initial genome-wide response to ECM stiffness and Rac. However, the mRNAs for two transcription factors (Klf2 and ATF3) were repressed by ECM stiffness with high statistical significance (Fig. 2E). Of those two, the repression of ATF3 mRNA was much more robustly reversed by inhibition of Rac activity with EHT1864 (Fig. 2F). Note that Figs. 2E-F also revealed a strong effect of ECM stiffness and Rac inhibition on the expression level of Dnajb1 mRNA. Though associated with fibrolamellar carcinoma as a fusion protein (Dinh *et al*., 2022), relatively little is known about Dnajb1 biology, and we therefore decided to focus our further studies on ATF3.

RT-qPCR confirmed that ATF3 mRNA levels were downregulated by culturing MEFs on a stiff substratum and strongly upregulated by treatment with EHT1864; the response to CT04 was not significantly different from the controls (Fig. 2G). Immunoblotting showed a similar repression of ATF3 protein abundance in response to ECM stiffness, and this effect was partially reversed when Rac-inhibited cells were cultured on a stiff ECM (Fig. 2G-I).

### Stiffness/Rac-mediated repression of ATF3 linked to the expression of cyclin D1

ATF3 belongs to the ATF/CREB family of transcription factors, has pleiotropic effects on cells, and is canonically induced in response to various stress signals (Hai *et al*., 1989; Hai and Hartman, 2001; Fan *et al*., 2002; Rohini *et al*., 2018; Ku and Cheng, 2020). Intriguingly, ATF3 has also been described as a repressor of cyclin D1 transcription, with direct binding to an inhibitory ATF3 site in the cyclin D1 promoter (James *et al*., 2006; Lu *et al*., 2006). As indicated above, cyclin D1 is a key regulator of progression through G1 phase, and we have previously shown that it is a major target of ECM stiffness-mediated signaling to the cell cycle (Klein *et al*.,2009; Bae *et al*., 2014). By plating FBS-stimulated MEFs on FN-coated hydrogels of increasing stiffness, we found that the levels of ATF3 mRNA (Fig. 3B) and cyclin D1 mRNA (Fig. 3C) both correlated with the level of Rac activity (Fig. 3A). In agreement with the results in Fig. 2, the correlation to Rac activity was inverse for ATF3 mRNA but direct for cyclin D1 mRNA.

**Figure 3.**
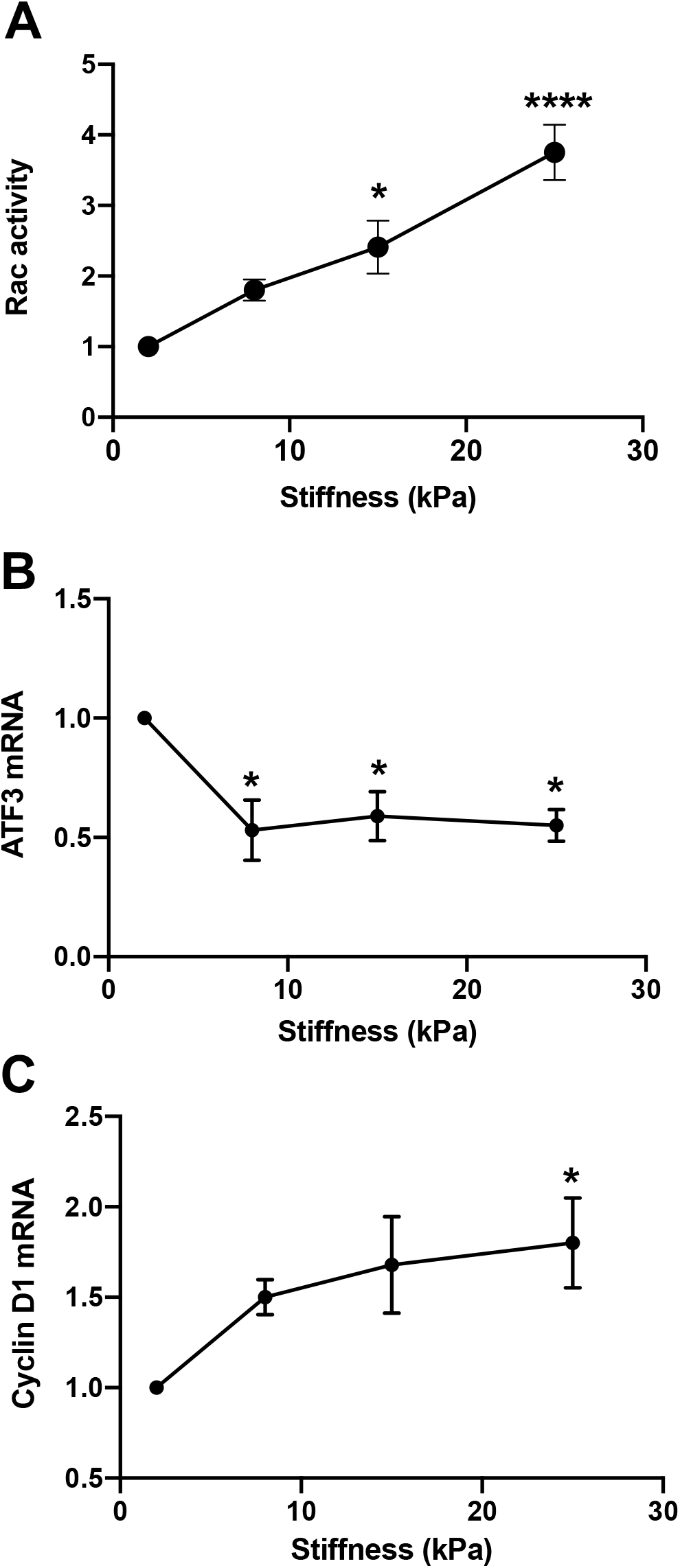
Dose-dependent effects of ECM stiffness on ATF3 and cyclin D1 mRNAs. **(A)** Serum-starved MEFs were seeded on FN-coated hydrogels of increasing stiffness (approximately 2, 8, 15, and 25 kPa) and incubated in DMEM-10% FBS for 1 h prior to cell lysis and analysis of Rac-GTP levels by G-LISA. Results show mean ± SEM (n=4). **(B-C)** Cells were treated as in panel A but incubated for 9 h prior to determination of ATF3 and cyclin D1 mRNA levels by RT-qPCR (n=3). Statistical significance for each panel was determined by ANOVA; asterisks show the results of Dunnett’s post-tests performed relative to the softest ECM (2-4 kPa) ECM.

We then compared the effect of Rac inhibition and activation on the expression of ATF3 and cyclin D1 mRNAs within the same lysates. RT-qPCR revealed a strong inverse correlation between the expression of ATF3 and cyclin D1 mRNAs in response to ECM stiffening (Fig. 4A; compare soft versus stiff). Moreover, both the stiffness-dependent repression of ATF3 mRNA and induction of cyclin D1 mRNA were reversed by Rac inhibition with EHT1864 (Fig. 4A; compare stiff ± EHT). Conversely, this inverse correlation was retained when we enforced Rac activity in MEFs on a soft ECM by ectopic expression of Rac^V12^: the level of ATF3 mRNA decreased while that of cyclin D1 mRNA increased (Fig. 4B). Importantly, this inverse relationship between stiffness-dependent ATF3 repression and cyclin D1 induction is causal because forced expression of ATF3 in cells on stiff hydrogels was sufficient to block the stiffness-dependent induction of cyclin D1 (Fig. 4C-D).

**Figure 4.**
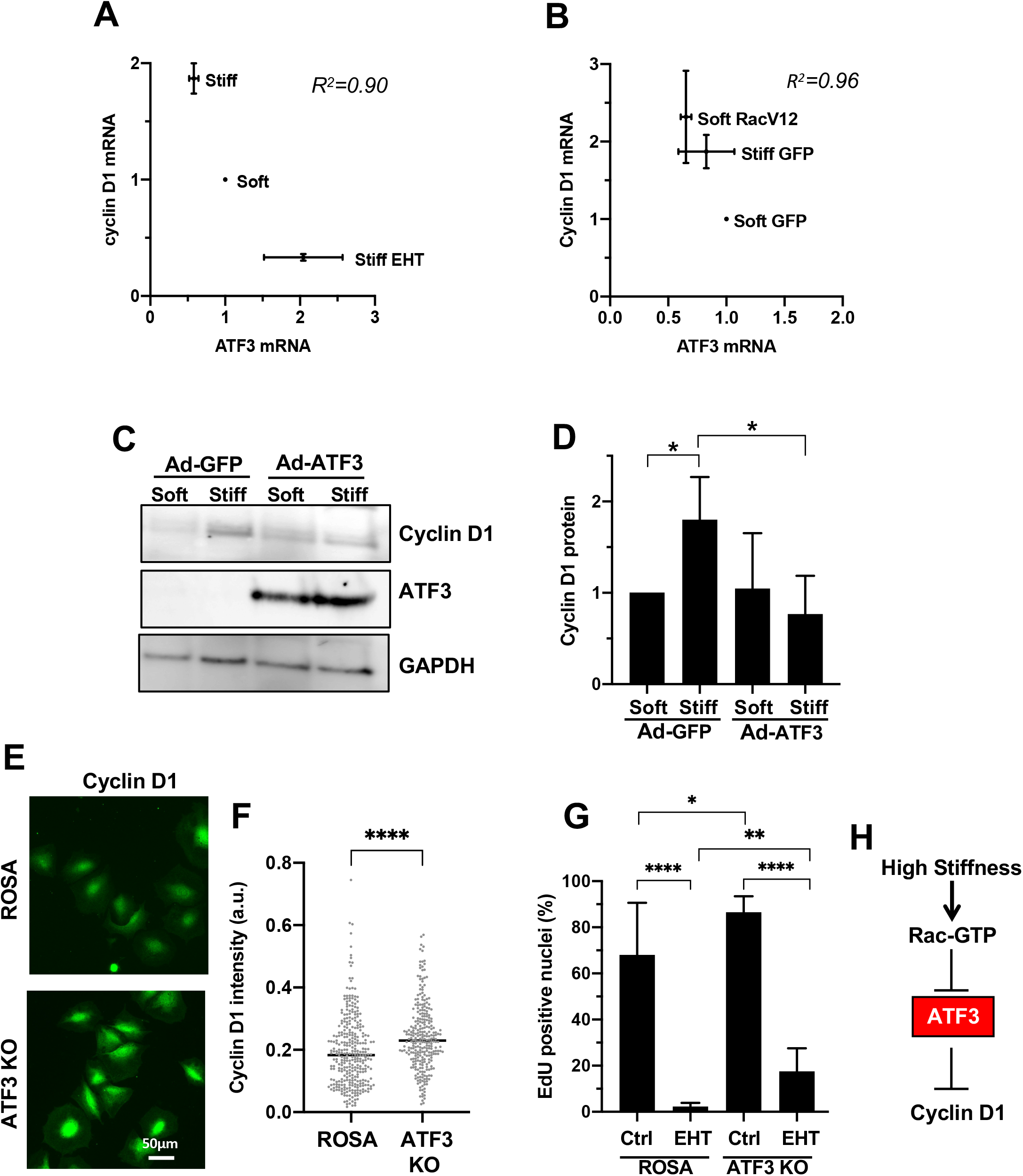
ATF3 repression linked to stiffness-dependent cyclin D1 expression. **(A)** Serum-starved MEFs were plated on soft or stiff FN-coated hydrogels in DMEM-10% FBS and treated with either vehicle (DMSO) or EHT1864 for 9 h. Cells were lysed, and the levels of both ATF3 mRNA and cyclin D1 mRNA were determined by RT-qPCR from the same lysates. Results show mean + SEM (n=3). R^2^ is the correlation coefficient. **(B)** MEFs were infected with adenoviruses encoding GFP (control) or Rac^V12^ and serum-starved. The cells were then cultured and analyzed by RT-qPCR as in panels A. Results show mean + SEM (n=3). **(C)** MEFs were infected with adenoviruses (Ad) encoding GFP (control) or ATF3. Cells were serum-starved and plated on FN-coated hydrogels for 15 h. The cells were analyzed by western blot for the level of cyclin D1 and ATF3 with GAPDH as the loading control. **(D)** Quantification of western blot results in C; the graph shows mean + SD (n=3). (**E)** Representative images of ROSA (control) and ATF3 KO MEFs that had been serum-starved and plated on stiff FN-coated hydrogels in DMEM-10% FBS for 9 h before fixation and immunostaining for cyclin D1. **(F)** Cyclin D1 staining intensity was quantified with Image J from ~ 300 cells per condition accrued from 2 independent experiments. The dot plot shows results for each cell and the mean. (**G)** S-phase entry was analyzed by EdU incorporation in 4 different ROSA (control) clones and 4 different ATF3 KO clones after serum-starvation incubation on stiff FN-coated hydrogels in DMEM-10% FBS for 24 h. Results show mean + SD; n=7 for the ROSA clones and 8 for the ATF3 KO clones. **(H)** Model showing how cyclin D1 is regulated by ECM stiffness and Rac through ATF3.

Crispr-Cas9 methodology was used to delete ATF3 from MEFs and determine the effect on cyclin D1 and S-phase entry. We generated several ATF3-deficient clones (hereafter called ATF3 KO cells) as well as several control clones in which MEFs were transfected with a ROSA26 rather than ATF3 gRNA (Table S5 and Fig. S2A-B; red arrowheads). Phenotypically, the lack of ATF3 had no apparent effect on the cells, nor did we detect large changes in cell area or f-actin intensity (Fig. S2B-D). However, serum-stimulated ATF3 KO MEFs on stiff hydrogels displayed an increase in cyclin D1 expression (Fig. 4E-F), and this increase was associated with an increase in S phase entry as judged by the percentage of EdU positive nuclei (Fig. 4G; compare columns 1 and 3). Rac inhibition with EHT1864 reduced S phase entry in both the control the ATF-depleted MEFs (Fig. 4F; compare bars 1 to 2 and 3 to 4), indicating (not surprisingly) that Rac activity drives cyclin D1 expression by pathways beyond ATF3. Note, however, that the extent of S phase inhibition by EHT1864 was 30-fold in the ROSA controls (from 68% to 2%) but only 5-fold in the ATF3 KO MEFs (from 87% to 18%). Altogether, these data demonstrate that a stiffness- and Rac-dependent repression of ATF3 drives the induction of cyclin D1 (Fig. 4H). This effect contributes not only to mechanosensitive cyclin D1 expression but also to the extent of S phase entry.

## DISCUSSION

The stiffness (or rigidity) of the ECM is transduced into intracellular stiffness and mechanosensitive signaling. Rac and Rho are important regulators of mechanical signaling, but the relative transcriptional response to Rac and Rho at the genome-wide level is not well understood. We have examined this question by characterizing the effects of these GTPases on the ECM stiffness-mediated transcriptome.

Given the established role of Rho on nuclear translocation of the MAL and YAP transcriptional co-activators, we were surprised to find that Rho inhibition had almost no effect on the early transcriptional response to ECM stiffness. Rather, we found that Rac played a major role, and controlled about 30-40% of all genes regulated by a stiff ECM, both positively and negatively. However, it is important to emphasize that the relative contributions of Rac and Rho to mechanosensitive transcription are likely to be time-dependent, and the transcriptional contribution of Rho may increase as cells attach, spread, and develop force in response to stiff substrata.

Our unbiased analysis identified ATF3 as a major target of the early transcriptional responses to both ECM stiffness and Rac activity. We found that ATF3 mRNA levels are repressed by a stiff ECM stiffness and active Rac, and that this repression is important for the stiffness- and Rac-dependent induction of cyclin D1. ATF3 binds to the cyclic AMP response element (CRE) (consensus: 5’-GTGACGT[AC][AG]-3’), in numerous promoters and can act as a transcriptional activator or repressor (Hai *et al*., 1989; Hai and Hartman, 2001; Fan *et al*., 2002; Rohini *et al*.,2018; Ku and Cheng, 2020). ATF3 homodimers can inhibit gene targets directly through an association with histone deacetylase 1, whereas ATF3-containing heterodimers can either positively or negatively regulate gene expression. ATF3 has also been described as both a positive and negative cell cycle regulator. While some reports suggest that ATF3 promotes cell proliferation (Allan *et al*., 2001; Perez *et al*., 2001), our results are in accordance with several other studies that have identified a negative role for ATF3 on cell cycle progression. For example, in addition to the inhibition of cyclin D1 gene expression as mentioned above, others have reported a repressive effect of ATF3 on cyclin A, cyclin E and Cdk2 levels, and ectopic expression of ATF3 suppresses cell cycle progression from G1 to S phase (Fan *et al*., 2002; James *et al*., 2006; Lu *et al*., 2006). Use of different cell types and experimental conditions likely affect the cellular response to alterations in ATF3 levels. However, our use of substrata of physiologically relevant stiffness is likely providing additional insight that is not attainable when cells are cultured on rigid (glass or plastic) surfaces. Studies in 3D may provide even more insight. How the positive and negative cell cycle effects of ATF3 relate to ECM rigidity, dimensionality, time, and dimerization are important, but likely complex, matters to understand. Coordination between the stiffness effects on ATF3 expression as shown here and nuclear mechanosensing (Dahl *et al*., 2008; Wang *et al*.,2009; Swift and Discher, 2014; Kirby and Lammerding, 2018) will also be of interest.

Increased tissue stiffness, typically a result of ECM remodeling, is seen in many pathological microenvironments, such as breast and pancreatic tumors, lung and liver fibrosis, and cardiovascular disease (Paszek *et al*., 2005; Duprez and Cohn, 2007; Levental *et al*., 2009; Liu *et al*., 2010; Keely, 2011; Kothapalli *et al*., 2012; Gehler *et al*., 2013; Tung *et al*., 2015; Piersma *et al*., 2020; Zhang *et al*., 2021; Maneshi *et al*., 2021) and has even been considered as a prognostic factor in cancer progression (Wei and Yang, 2016; Reid *et al*., 2017). Our data indicate that ATF3 is an early and major transcriptional target of increased ECM stiffness. Since ATF3 has widespread effects on cells, its downregulation by pathologically stiff microenvironments may be an early event that amplifies transcriptional misregulation in mechanosensitive pathologies.

## METHODS

### Cell culture, pharmacologic inhibition, ectopic expression, and GTPase activity assays

Spontaneous immortalized MEFs were cultured in DMEM-10% FBS at 37°C with 10% CO_2_. For serum-starvation, near confluent monolayers were incubated for 48 h with DMEM-1 mg/ml heat-inactivated fatty-acid free BSA. The cells were cultured on polyacrylamide hydrogels, prepared as previously described (Klein *et al*., 2007; Cretu *et al*., 2010) on 12-mm, 25-mm or 40-mm coverslips and coated with FN (#341631, EMD) at 5 μg/ml. Hydrogel stiffness ranged from 2–4 kPa (low) to 20–24 (high) kPa. The coverslips were collected, and the cells were fixed for immunofluorescence (12-mm coverslips), or extracted for RNA (25-mm coverslips) or protein (40-mm coverslips).

#### Acute inhibition of Rac or Rho

MEFs were cultured in DMEM-10% FBS to 70-80% confluency and then serum-starved in DMEM-1 mg/ml BSA. After 48 h, the cells were trypsinized, resuspended in DMEM-1%BSA and incubated in suspension for 30 min at 37°C in 10% CO_2_ and 2 μg/ml CT04 (#CT04-A, Cytoskeleton or 10 μM EHT1864 (#3872, Tocris). The vehicle control included the corresponding dilution of DMSO as needed. These pretreated cells were collected by gentle centrifuged, resuspended in DMEM-10% FBS, and plated on FN-coated hydrogels in the continued presence of inhibitor for 1 h.

#### Ectopic expression

MEFs were cultured at 70-80% confluency and infected with adenoviruses encoding Rac^V12^ (kind gift of Chris Chen, Boston University), Ad-GFP-h-ATF3 (Vector Biolabs), or GFP largely as described (Klein *et al*., 2007; Bae *et al*., 2014). The adenovirus dilutions used resulted in ~80% cell infection as judged by the GFP signal. The infected cells were washed with DMEM-1 mg/ml BSA, serum-starved, trypsinized, and plated on FN-coated polyacrylamide hydrogels as described above for 9 h for analysis by RT-qPCR and western blotting.

#### Rho family GTPases activity assay

Cells were treated with EHT1864 and CT04 as described above were collected, and Rac or Rho activity was assessed in duplicate using the G-LISA small G-protein activation assay kit (BK128, BK124, Cytoskeleton) according to the manufacturer’s directions. Briefly, serum-starved MEFs were replated on fibronectin-coated low- and high-stiffness hydrogels and stimulated with 10% FBS for 1. Total cell lysates were prepared with icecold lysis buffer and the protein concentration was measured by Coomassie binding (Bio-Rad). Equal amounts of protein were added to each well of a G-LISA plate and incubated for 30 min at 4°C. The bound Rac was detected by adding an anti-Rac primary antibody and a secondary HRP-labeled antibody to each well. Finally, HRP detection reagents were added, and the resulting colorimetric reaction was quantified by measuring absorbance at 490 nm in a microplate spectrophotometer.

### Atomic force microscopy (AFM)

Intracellular stiffness was measured by plating cells for 1 h on 18-mm soft- or stiff FN-coated polyacrylamide hydrogels with 10% FBS. The intracellular stiffness of single adherent cells was measured using a DAFM-2X Bioscope (Veeco) mounted on an Axiovert 100 microscope (Zeiss) in contact mode. Cells were indented against a standard silicon nitride cantilever (spring constant = 0.06 N/m) with a conical tip (40 nm in diameter). The elastic modulus (stiffness) was calculated by fitting the first 600 nm of tip deflection from the horizontal with the Hertz model for a cone. The tip was placed near the edge of the cell to measure intracellular stiffness. Measurements were taken for 7–10 cells per condition, and the data were analyzed using a custom MATLAB script generously provided by Paul Janmey (University of Pennsylvania).

### Fluorescence microscopy and quantification of f-actin intensity

Hydrogels were washed with PBS, and the cells were fixed in 3.7% formaldehyde (room temperature for 15 min) washed 3 times with PBS, and permeabilized with 0.4% Triton X-100 in PBS for 10 min, washed once with PBS, and blocked in 2% BSA and 0.2% Triton X-100 (30 min at room temperature). Primary antibodies to ATF3 (sc-188, Santa Cruz) and cyclin D1 (sc-450, Santa Cruz) were diluted at 100-fold in 2% BSA, 0.2% Triton X-100 and incubated with the cells for 1 h. The cells were then washed 3 times in PBS, 2%, BSA/0.2% Triton X-100, incubated with secondary antibody (diluted 100-fold) for 1 h at room temperature. The immunostained cells were then washed twice, and the coverslips were mounted using DAPI fluoromount G (0100-20, SouthernBiotech). To quantify F-actin signal intensity, fixed cells that had been stained for 1 h with Alexa Fluor™ 594 Phalloidin (A12381, ThermoFisher) in 2% BSA and 0.2% Triton X-100 were washed 3 times in PBS/0.2% Triton X-100 and mounted as described above.

Quantification of cell area was performed using ImageJ: a threshold was set to cover the whole cell surface, holes were filled using Process>Binary>Fill Holes commands, and then the individual cell areas were measured (Dang *et al*., 2013). The phalloidin signal for each cell was measured, set on an arbitrary threshold based on the DMSO control cell intensity. F-actin intensity was then normalized to cell area for each cell.

### RNASeq and bioinformatic analysis

Quadruplicate samples were generated for MEFs cultured in 4 different conditions: i) soft hydrogels, ii) stiff hydrogels, iii) stiff hydrogels with EHT1864, and iv) stiff hydrogels with CT04 as described above for “Acute Inhibition of Rac or Rho.” Total RNA was extracted using Trizol reagent (Invitrogen), further purified with an RNeasy kit (74106, Qiagen), and prepared for RNA sequencing using TruSeq RNA Stranded mRNA (Illumina) and PE100. Sequencing coverage was ~40×10^6^ reads. Salmon (https://combine-lab.github.io/salmon/) was used to count the data against the transcriptome defined in Gencodev M28, which was built on the genome GRCm39. Several Bioconductor packages (https://bioconductor.org) were used for subsequent steps. The transcriptome count data was annotated and summarized to the gene level with tximeta and further annotated with biomaRt. A Principal Component Analysis was performed with PCAtools, and this led to the exclusion of 1 of the 4 replicates of MEFs cultured on a soft hydrogel. Raw feature counts of the remaining 15 samples were normalized and analyzed for differential expression using DESeq2. Venn diagrams were generated from the DESeq2 output based on cut-offs of 0.32 log_2_ (approximately >1.25 log_10_) fold-change (positive or negative), <0.05 adjusted p-value, and >500 for base mean intensity. The genes regulated by ECM and Rac were compared to GO gene lists for transcription factors, transcription co-regulators and histone modifiers (GO terms 0003700, 0003712, and 0016570, respectively).

### CRISPR/Cas9-mediated deletion of ATF3

The CRISPR sgRNA to mouse ATF3 (CCAGCGCAGAGGACAUCCGA) and ROSA26 (GAACAUAAAUGGCAACAUCU) were obtained from Synthego Corporation. The sgRNAs were diluted to 30 μM and Cas9 Nuclease to 20 μM. MEFs were seeded in 12-well plates and grown to 70-80% confluency. Cells were transfected with RNPs (ribonucleoprotein) complexes using a NEON electroporation system (Invitrogen) and an RNP Complexes ratio of 9:1. Cells were electroporated with the following parameters: 1 pulse at 1350 V and 30 ms, then incubated in DMEM-10% FBS for 2-3 days. For clonal expansion, each well was trypsinized and diluted such that an average of 0.5 cells were plated per well in 96-well plates. Cells were cultured until they reach ~70% confluency and then subjected to Sanger sequencing. ROSA26 primers were ACATTTGGTCCTGCTTGAACA (forward) and ACATTTGGTCCTGCTTGAACA (reverse). ATF3 primers (for gRNA-1) were GTAGGCTGTCAGACCCCATG (forward) and GGTGCACACTATACCTGCTC (reverse). Sanger sequencing data were uploaded to the ICE CRISPR Analysis Tool (Synthego) to assess the ATF3 editing efficiency for each clone. Clones with a predicted knockout efficiency >95% were analyzed further by Western Blotting.

### Statistical analysis

Statistical significance was determined using Prism (Graph-Pad) software. Graphs show means + SD unless the independent experiments generated means, in which case the error bars show SEM. T-tests were used to compare data unless noted otherwise in a figure legend. T-tests were 2-tailed unless testing for an effect in a specific direction. Unless specified in the figure legend, statistical significance was determined by t-tests with *p<0.05, **p<0.01, ***p<0.001, and ****p<0.0001. The reciprocal relationship between ATF3 and cyclin D1 was assessed using the correlation coefficient, R^2^.

## ACKNOWLEDGEMENTS

We John Tobias (University of Pennsylvania) for assistance with the bioinformatic analysis This work was supported by NIH grant HL137232 to RKA, by the Center for Engineering MechanoBiology, a National Science Foundation Science and Technology Center under grant agreement CMMI 15-48571, and by EMBO post-doctoral EMBO ALTF 1082-2016 to ID. YB was supported by American Heart Association Career Development Award 18CDA34080415.

