## supplemental methods-figs-tables for "Rac rather than Rho drives the early transcriptional response to extracellular matrix stiffness and mediates repression of ATF3"

**RT-qPCR.** Total RNA was extracted and purified from MEFs using the Quick-RNA Miniprep Plus Kit (Zymo research). cDNAs were prepared using equal amount of total RNA diluted into TaqMan® reverse transcription master mix (ThermoFisher) and then processed according to the manufacturer's instructions. Gene expression was quantified by RT-qPCR performed in duplicate using TaqMan® Universal PCR master mix, ~20 ng of total RNA for reverse transcription (RT), and 5 ng of RT product in the qPCR. Taqman assays (ThermoFisher) were used for ATF3 (Mm00476033_m1) and CCND1 (Mm00432359_m1) The primer probe set for 18S rRNA has been described (Klein *et al.*, 2007). The level of mRNA expression for each gene was determined by the ddCt method and plotted relative to 18S rRNA.

**Immunoblotting**. MEFs on hydrogels were collected in RIPA buffer (50mM Hepes, 150mM NaCl, 1% NP-40, 0.5% deoxycholate, and 0.1% SDS) containing protease inhibitors (Cell Signaling; 5872S). Cell lysates were centrifuged to remove nuclei and debris, Supernatants were collected, and their protein concentrations were determined using a Bradford protein assay (Bio-Rad Laboratories; 5000006). Equal amounts of proteins were diluted in LDS Sample Buffer (Invitrogen™) and fractionated on 12% SurePage precast gels (Genscript). In some experiments, the cells were extracted in 5 x SDS sample buffer as described (Bae *et al.*, 2014). Proteins were transferred onto nitrocellulose or polyvinylidene difluoride (PVDF) membranes. The membranes saturated with 5% BSA in 1 x TBS (20 mM Tris, pH 7.5, 150 mM NaCl) with 0.1% Tween-20 and probed with primary antibodies to ATF3 (Santa Cruz, sc-188 or Novus Biologicals, NBP1-85816), cyclin D1 (Santa Cruz, sc-20044) or GAPDH (loading control; ThermoFisher, MA5-15738) and secondary antibodies ECL anti-rabbit Hrp (GE Healthcare, 3144,) and EC anti-rabbit HRP (GE Healthcare, 3143). Antibodies were diluted in the same TBS buffer, and signals were detected by enhanced chemiluminescence with an ImageQuant Las4000.

**EdU incorporation assay**. Cells were starved in DMEM-1 mg/ml BSA for 48 h, trypsinized, and incubated for 24 h on hydrogels in DMEM-10% FBS and 10 µM 5-ethynyl-2′-deoxyuridine (EdU; Invitrogen). EdU was visualized using the Click-iT EdU Imaging Kit (Invitrogen) according to the manufacturer's instructions. Coverslips were mounted onto glass microscope slides using DAPI fluoromount G (SouthernBiotech, 0100-20,) for manual counting of DAPI-stained and EdU-positive nuclei.

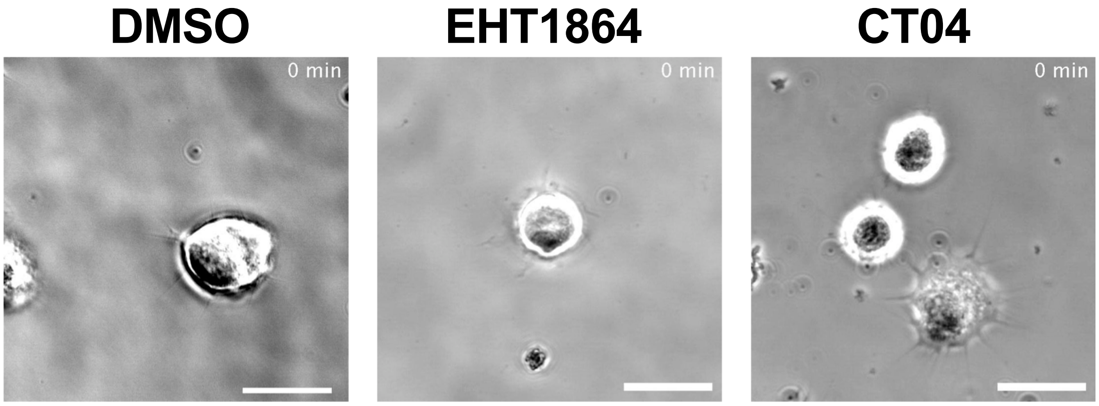


**Figure S1.** MEFs were cultured in DMEM-10% FBS to 70-80% confluency, trypsinized, and pre-incubated in suspension for 30 min in DMEM with 1 mg/ml BSA and either vehicle (DMSO), EHT1864 or CT04. The cells were then plated on FN-coated stiff hydrogels in DMEM-10% FBS in the continued presence of the inhibitors as described in Methods. The cells were imaged every 15 sec for 15 min using the 40X objective of a Zeiss Axio Observer 7 inverted microscope. Scale bar = 30 µm. Movies were selected to show typical responses; n~100 cells accrued from 2 independent experiments.

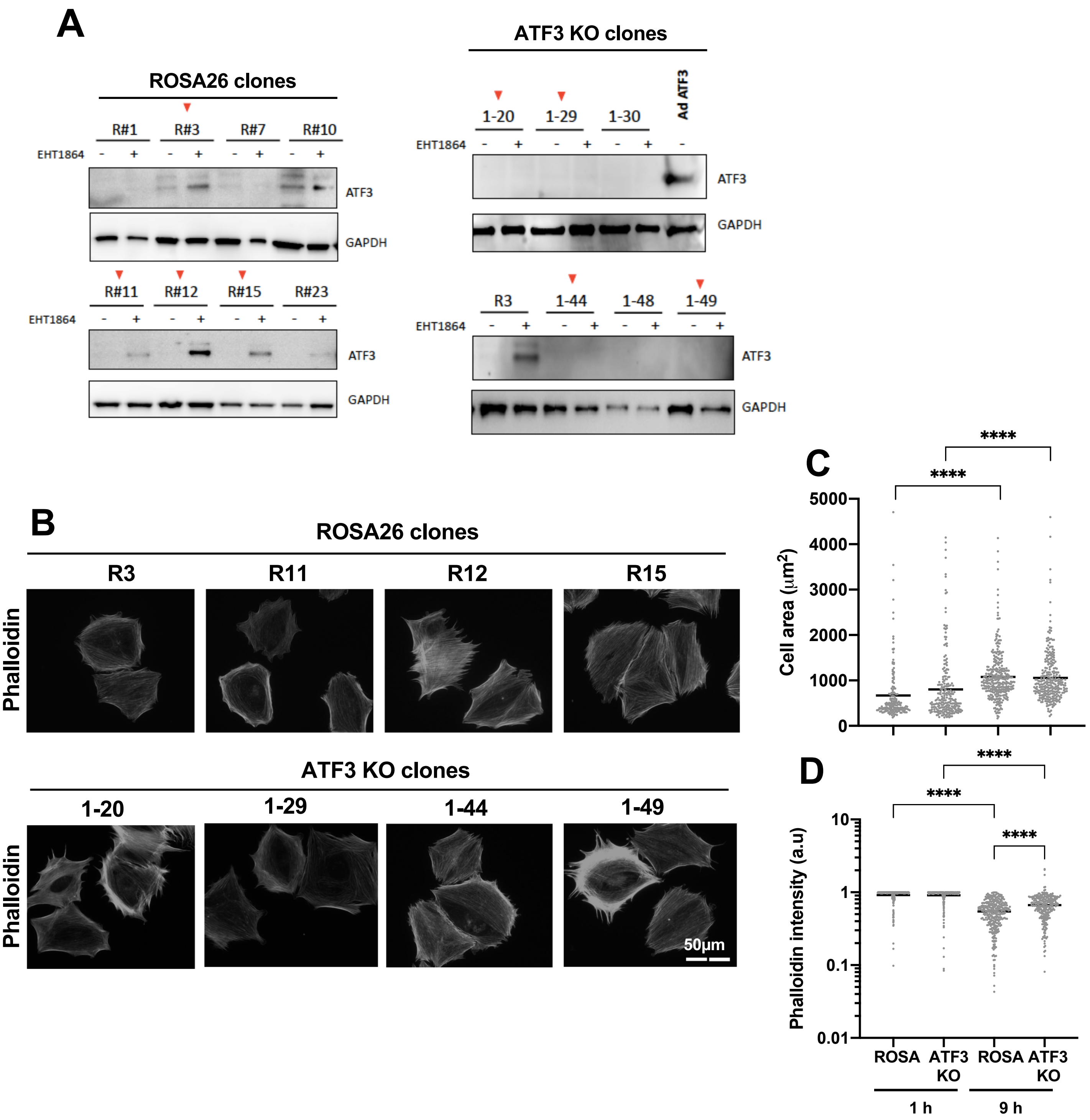


**Figure S2. Identification and initial characterization of ATF3 knock-out (KO) MEFs on stiff hydrogels.** (**A**) Several putative ATF3 deletion clones, including those identified by the ICE Crispr Analysis Tool (Table S5), and a similar number of ROSA26 (control) clones were serum-starved, pre-treated with 10 mM EHT1864 in suspension, and plated on FN-coated tissue culture dishes with 10% FBS-DMEM in the continued presence of EHT1864 for 5 h before determination of ATF3 protein levels by western blot. GAPDH was the loading control. Red arrowheads show the clones selected for further analysis. **(B)** Images of control and ATF3 KO MEFs that had been serum-starved and plated on stiff FN-coated hydrogels in DMEM with 10% FBS for 9 h before fixation and staining with Alexa Fluor™ 594 Phalloidin. **(C-D)** Quantitative analysis of cell area and phalloidin intensity in the ROSA26 control (R3, R11, R12, R15) and ATF3 KO clones (1-20, 1-29, 1-44, 1-49) using Image J. The graphs show ~200-300 cells analyzed per condition accrued from 2 independent experiments.

**Table S1. Gene lists for the Venn diagrams shown in Figure 2A.**

**I) Stiffness-stimulated and EHT1864-inhibited** (157 genes)

1810013L24Rik
Adamts5
Aebp2
Ahctf1
Ahnak
Akap11
Alg10b
Amer1
Ankhd1
Ankrd17
Ankrd50
Apaf1
Arc
Arhgap35
Arl5a
Atg2b
Birc6
Bmp2k
Bmpr2
Bpnt2
Bptf
Btaf1
Btbd7
Cbfa2t2
Ccnt1
Cdk6
Cert1
Clasp2
Cltc
Cmtm4
Cnot6
Crybg1
Csnk1g1
Dicer1
Dido1
Dip2b
Dmxl1
Dnm3os
Dock5
Dock9
Dr1
Dyrk1a
Elk4
Errfi1
Exoc5
F2r
Fam117b
Fem1b
Fmr1
Garre1
Ggnbp2
Gm14636
Gm37899
Gtf2a1
Hdac4
Hectd1
Kdm5b
Kif13b
Kifap3
Klhl11
Kmt2c
Lats1
Lats2
Lgr4
Lnpep
Lrrc58
Mast4
Mdn1
Med13
Med14
Mical3
Mindy2
Mob1b
Mrtfb
mt-Nd5
Mycbp2
Naa15
Naa50
Nf1
Npat
Nufip2
Nup153
Nup98
Pafah1b1
Pak2
Pank3
Parg
Pdzd8
Peak1
Phc3
Pigm
Ppp1r15b
Prkdc
Psme4
Purb
Rab1a
Ranbp2
Rap1b
Rap2c
Rasa1
Rbbp6
Rc3h2
Ric1
Rictor
Rlf
Rlim
Rmnd5a
Rnf111
Rnf169
Rnf213
Runx1
Sacs
Secisbp2l
Shroom4
Six4
Smg7
Sos1
Sos2
Spred1
Stard9
Sypl
Tanc1
Tasor
Tasor2
Tbc1d8b
Tent4b
Tjp1
Tmem263
Top1
Tpp2
Trib1
Tshz1
Ttc14
Ubr5
Uhmk1
Uhrf1bp1l
Urb2
Usp14
Usp24
Usp9x
Vcpip1
Washc4
Wdr26
Xpo4
Xrn1
Yeats2
Ythdf3
Zbed6
Zbtb1
Zfc3h1
Zfp267
Zfp275
Zfp281
Zfp948
Zfpm2
Zhx3
Zswim6

**II) Stiffness-stimulated and both EHT1864- and CT04-inhibited** (1 gene)

Zfp646

**Table S2. Gene lists for the Venn diagrams shown in Figure 2B.**

**I) Stiffness-inhibited and EHT1864-stimulated** (109 genes)

1110038B12Rik
2410006H16Rik
4933434E20Rik
A430005L14Rik
Aimp1
Arfip2
Atf4
Atp6v0b
Atpif1
Bloc1s1
Bola2
Cinp
Cnpy2
Commd1
Cox7a2l
Cst3
Cstf2
Ctsh
Czib
Ddit3
Eef1b2
Erp29
Fau
Ftl1
Gadd45a
Gm10275
Gm11808
Gm12481
Gm14494
Gm15500
Gm2000
Gm26532
Gm4332
Gm6969
Gm7536
Gm8210
Gnptg
Grb2
H1f2
H2ac19
H2bc4
Hoxb2
Ift22
Igbp1
Igfbp2
Jun
Jund
Mri1
Mrpl19
Mrpl21
Mrpl30
Mrpl52
Mt2
Mydgf
Ndufa3
Ndufa6
Ndufa7
Ndufb5
Neurl4
Ost4
Paxx
Pcif1
Rpl11
Rpl12
Rpl13
Rpl14
Rpl15-ps2
Rpl15-ps6
Rpl18
Rpl18a
Rpl21
Rpl22
Rpl23a
Rpl24
Rpl31-ps8
Rpl34
Rpl35
Rpl37
Rpl4
Rpl6
Rpl8
Rps12
Rps13-ps1
Rps17
Rps18
Rps19
Rps20
Rps21
Rps23-ps1
Rps25
Rps27a
Rps27rt
Rps3
Rps5
Rps6
Rps7
Rps7-ps3
Rps8
Rps9
Snhg16
Stoml2
Tmed1
Tmem208
Tmem242
Tpt1-ps3
Uqcc2
Uqcr10
Washc3
Zfas1

**II) Stiffness-inhibited and both EHT1864- and CT04-stimulated** (14 genes)

Atf3
Dnaja4
Dnajb1
Gm12338
Hspa1a*
Hspa1b
Hspb1
Klf2
Mrps33
Nedd8
Rhob
Rpl31
Rpl36a-ps2
Serf1

*not pursued as its calculated adjusted p-value in DESeq2 was undefined

**Table S3. Gene lists for the Venn diagrams shown in Figures 2C-D.**

**I) Stiffness-stimulated and EHT1864-inhibited**

*Transcription factors & co-regulators (23 genes)*

| | | |
|---|---|---|
| Aebp2 | Npat | Zbtb1 |
| Btaf1 | Nup98 | Zfp267 |
| Cbfa2t2 | Purb | Zfp275 |
| Cnot6 | Rap2c | Zfp281 |
| Dyrk1a | Runx1 | Zfp948 |
| Med13 | Six4 | Zfpm2 |
| Med14 | Tshz1 | Zhx3 |
| Mrtfb | Zbed6 | |

II) *Histone modifications* (6 genes)

Fmr1
Ubr5
Parg
Yeats2
Naa50
Dr1

III) *Histone modifications and Transcription factors and co-regulators (5 genes)*

Elk4
Hdac4
Kdm5b
Kmt2c
Rlf

**II) Stiffness-inhibited and EHT1864-stimulated**

*Transcription factors & coregulators (9 genes)*

Atf3
Atf4
Ddit3
Dnajb1
Hoxb2
Hspa1a
Jun
Jund
Klf2

*Histone modifications (1 gene)*

H1f2

**Table S4. Gene list, fold changes (FC) and adjusted p-values (padj) for the graphs shown in Figures 2E-F.**

| Gene | FC stiff | padj stiff | FC EHT | padj EHT |
|---|---|---|---|---|
| Aebp2 | 0.362360035 | 0.039872997 | -0.430363934 | 0.003114619 |
| Atf3 | -0.842902142 | 1.24811E-11 | 2.145058258 | 1.93545E-98 |
| Atf4 | -0.37550875 | 0.00280697 | 1.094335394 | 1.56274E-34 |
| Btaf1 | 0.464672293 | 0.008971385 | -0.324555895 | 0.04343359 |
| Cbfa2t2 | 0.41488484 | 0.011717977 | -0.339958504 | 0.018569795 |
| Cnot6 | 0.503573055 | 0.013761436 | -0.447533286 | 0.011382425 |
| Ddit3 | -0.455580575 | 0.017797817 | 1.232648422 | 2.23548E-19 |
| Dnajb1 | -0.823217294 | 3.24098E-10 | 3.039256505 | 2.3813E-185 |
| Dr1 | 0.478122321 | 0.013496317 | -0.416349259 | 0.01323619 |
| Dyrk1a | 0.40944526 | 0.004902845 | -0.42730384 | 0.000374685 |
| Elk4 | 0.441802107 | 0.008318563 | -0.518755581 | 0.000141077 |
| Fmr1 | 0.64383477 | 0.002564592 | -0.414247267 | 0.032879518 |
| H1f2 | -0.599990161 | 0.010656195 | 0.965007028 | 7.64479E-08 |
| Hdac4 | 0.357327008 | 0.016954218 | -0.582158879 | 5.14256E-07 |
| Hoxb2 | -0.396567327 | 0.014372579 | 0.437993522 | 0.001075301 |
| Jun | -0.537149245 | 0.005858218 | 1.801131885 | 6.29779E-40 |
| Jund | -0.664710836 | 3.55088E-07 | 0.881502801 | 2.63829E-16 |
| Kdm5b | 0.394928769 | 0.009067506 | -0.444544135 | 0.000317503 |
| Klf2 | -1.32456878 | 7.17855E-14 | 1.096519866 | 8.85946E-12 |
| Kmt2c | 0.605275299 | 0.041782327 | -0.595477572 | 0.019929676 |
| Med13 | 0.937533168 | 0.001556194 | -0.584996931 | 0.030493793 |
| Med14 | 0.425412702 | 0.04245209 | -0.52651539 | 0.002082577 |
| Mrtfb | 0.47663481 | 0.013024444 | -0.414162701 | 0.012859084 |
| Naa50 | 0.654766699 | 5.21957E-05 | -0.395677938 | 0.007332563 |
| Npat | 0.423277831 | 0.018719279 | -0.422098456 | 0.005748077 |
| Nup98 | 0.441426877 | 0.017037315 | -0.328370145 | 0.049235075 |
| Parg | 0.45320743 | 0.027580408 | -0.410129915 | 0.022023721 |
| Purb | 0.851574605 | 0.001415271 | -0.73580509 | 0.001118295 |
| Rap2c | 0.52841838 | 0.000446182 | -0.432019665 | 0.000695973 |
| Rlf | 0.41422215 | 0.006242845 | -0.636060685 | 5.16445E-08 |
| Runx1 | 0.692994542 | 0.001947058 | -0.952609268 | 5.12789E-08 |
| Six4 | 0.628202539 | 0.002009872 | -0.998712042 | 1.47086E-10 |
| Tshz1 | 0.445936334 | 0.039202763 | -0.556967227 | 0.001604112 |
| Ubr5 | 0.538590616 | 0.022150642 | -0.459121996 | 0.026967911 |
| Yeats2 | 0.408365022 | 0.012058095 | -0.389266858 | 0.004684534 |
| Zbed6 | 0.714189321 | 0.030122649 | -0.723758197 | 0.00948818 |
| Zbtb1 | 0.456335088 | 0.004845731 | -0.70166333 | 2.05241E-08 |
| Zfp267 | 0.354369365 | 0.021576453 | -0.391180118 | 0.002202342 |
| Zfp275 | 0.414505347 | 0.001832134 | -0.450659831 | 3.11125E-05 |
| Zfp281 | 0.448880655 | 0.022560895 | -0.94108134 | 1.93456E-10 |
| Zfp948 | 0.715492521 | 0.002439758 | -0.628019391 | 0.001681668 |
| Zfpm2 | 0.506331146 | 0.029174177 | -0.524693627 | 0.00734088 |
| Zhx3 | 0.511305749 | 0.000295488 | -0.381625136 | 0.001734441 |

| Clone ID | KO-Score | R-squared |
|---|---|---|
| 1-5 | 95 | 0.95 |
| 1-20 | 94 | 0.94 |
| 1-29 | 94 | 0.94 |
| 1-48 | 94 | 0.94 |
| 1-44 | 94 | 0.94 |
| 1-49 | 95 | 0.95 |

**Table S5. Identification of putative ATF3-depleted MEF clones**. MEF clones from the Crispr transfection were lysed in QuickExtract™ DNA Extraction buffer (Lucigen). Genomic DNA was extracted by two cycles of vortexing (15 sec) and heating (first at 68°C for 15 min and then at 95°C for 10 min). The DNA was subjected to PCR amplification, the purified PCR products were sequenced using a reverse ATF3 primer for gRNA-1 (GGTGCACACTATACCTGCTC), and the sequences were analyzed using the ICE CRISPR Analysis Tool (Synthego). A Knockout (KO) Score was generated for each clone. The KO score represents the proportion of cells that have either a frameshift or 21+ bp indel. We selected several clones with KO scores >90 for further analysis.
